# GRASP55 Safeguards Proper Lysosome Function by Controlling Sorting of Lysosomal Enzymes at the Golgi

**DOI:** 10.1101/2024.12.10.627846

**Authors:** Julian Nüchel, Maryam Omidi, Stephanie A. Fernandes, Marina Tauber, Sandra Pohl, Markus Plomann, Constantinos Demetriades

## Abstract

Lysosomes are multifunctional organelles that play important roles in cellular recycling, signaling, and homeostasis, relying on precise trafficking and activation of lysosomal enzymes. While the Golgi apparatus plays a central role in lysosomal enzyme sorting, the mechanisms linking Golgi function to lysosomal activity remain incompletely understood. Here, we identify the Golgi-resident protein GRASP55, but not its paralog GRASP65, as a key regulator of lysosome function. More specifically, we demonstrate that loss of GRASP55 expression leads to missorting and secretion of lysosomal enzymes, lysosomal dysfunction and bloating. GRASP55 deficiency also disrupts lysosomal mTORC1 signaling, reducing the phosphorylation of its lysosomal substrates, TFEB and TFE3, while sparing its non-lysosomal targets. Mechanistically, GRASP55 interacts with GNPTAB, a critical enzyme required for mannose 6-phosphate (M6P) tagging of lysosomal enzymes, and is necessary for its correct trafficking and stability. These findings reveal an essential role for GRASP55 in Golgi-lysosome communication and lysosomal enzyme trafficking, and suggest that *GRASP55/GORASP2* may act as a susceptibility gene for lysosomal storage disorder (LSD)-like conditions. Overall, this work underscores the importance of Golgi-mediated protein sorting in lysosome function and lysosomal mTORC1 signaling, and provides insights into the molecular basis of LSD-related pathologies.

## Introduction

Lysosomes are the main degradative organelles in cells. By breaking down their content, lysosomes recycle material on a continuous basis, thereby providing cells with fresh building blocks to support anabolic functions ^1–4^. Work over the last 15 years has expanded this view further, showing that the lysosomal surface also serves as a signaling platform for a number of key signaling molecules, like the growth- and metabolism-related mTOR (mechanistic/mammalian Target of Rapamycin) kinase ^5–8^, or its upstream negative regulator, the tumor suppressor TSC (Tuberous Sclerosis Complex) protein complex ^9–13^, both of which are regulated by nutrient and stress signals at this subcellular location to coordinate anabolic and catabolic processes in cells ^14–16^. At the same time, lysosomes remove insoluble aggregates and damaged proteins or even organelles, whose accumulation can cause severe cell damage, and is a hallmark of neurological disorders such as Alzheimer’s and Parkinson’s ^17–19^, thus underscoring the importance of proper lysosomal function for human disease and ageing ^3,4^.

Lysosomal cargo is degraded by soluble lysosomal enzymes that break down a variety of structurally diverse compounds, such as proteins, lipids, sugars, nucleic acids, and other macromolecules. The directed vesicle-mediated transport of newly synthesized lysosomal enzymes to these organelles involves a complex system of sorting signals and recognition proteins, a process that takes place in the Golgi apparatus, the main processing and distribution center for cargo proteins in cells ^4,20–22^. One such well-described signal is the modification of specific mannose residues on lysosomal enzymes that reach the Golgi from the endoplasmic reticulum (ER) and the ER-Golgi intermediate compartment (ERGIC) by the addition of phosphate groups to generate mannose 6-phosphate (M6P) ^23–28^. The key enzyme in the formation of M6P residues is the heterohexameric (α_2_β_2_γ_2_) GlcNAc-1-phosphotransferase (GNPT) complex. The *GNPTAB* gene encodes the transmembrane α/β precursor of GNPT, whereas *GNPTG* encodes the soluble γ subunit ^29^. After assembly of the GNPT subunits in the ER, the inactive complex is transported to the *cis*-Golgi and the α/β precursor (here referred to as GNPTAB) is proteolytically processed by site-1 protease (S1P) into the enzymatically active α and β subunits ^30–32^ (here referred to as GNPTA and GNPTB, respectively). The M6P residues on lysosomal proteins are then recognized by M6P receptors (MPRs) in specialized *trans*-Golgi subdomains, which facilitate their transportation to late endosomes and lysosomes ^22,24,25,33–35^. Lysosomal enzyme activity requires the acidic environment inside the lysosomal lumen that is established by the V-ATPase (vacuolar H^+^-ATPase) and other transmembrane ion channels ^36^. Besides ensuring that lysosomal enzymes are only active inside lysosomes, the acidic environment of the lumen plays an important role also for their delivery to these organelles, as MPRs bind M6P-tagged proteins at pH 6.7 in the *trans*-Golgi and release it in the endosomes at pH 6 due to changes in their affinity and conformation^34,37^.

Pathogenic variants in many lysosomal enzymes or proteins participating in their maturation and trafficking have catastrophic effects at the cellular and organismal levels, and cause lysosomal storage disorders (LSDs) in humans. More than 70 genetically distinct types of LSDs have been reported to date ^4,22,38,39^. Although LSDs are individually rare, collectively, they are found at relatively high incidence, with more than 1 in 5000 live births affected in the general population ^4,39,40^. While the symptomatology varies a lot between disorders in the LSD spectrum, most diseases affect the central nervous system, while some manifest with musculoskeletal system defects, impaired growth, and reduced life span ^4,39^. Whereas the vast majority of LSDs are caused by pathogenic variants in genes that code for soluble lysosomal enzymes or lysosomal membrane proteins, genetic defects in *GNPTAB*, encoding the catalytic subunits of the Golgi-resident GNPT complex, result in mistargeting of lysosomal enzymes and the severe human disease mucolipidosis type II (ML-II, formerly known as I-cell disease) ^41,42^. Cells from these patients are characterized by accumulation of non-degraded macromolecules, lysosomal bloating, and missorting of lysosomal enzymes that are secreted to the extracellular space as inactive proenzymes, instead of being delivered to lysosomes. Therefore, a key diagnostic feature in ML-II is the increased levels of lysosomal enzymes in the circulation ^29,42^. Highlighting its key role in maintaining cellular physiology, perturbation of Golgi function is also linked to cancer growth, invasion and metastasis, in addition to its involvement in the etiology of LSDs (reviewed in ^43^).

Unconventional protein secretion (UPS) is an alternative secretory route via which specific cargo proteins reach the cell surface or the extracellular space ^44^. In contrast to bulk secretory pathways, UPS is induced upon nutrient starvation or stress, and is, therefore, part of the cellular adaptation to stress. A central regulator of UPS is the Golgi-resident protein GRASP55 (Golgi re-assembly and stacking protein 55; also referred to as GORASP2) ^45–47^. Besides controlling UPS, together with the closely-related GRASP65 protein, GRASP55 was originally described to be involved in the assembly and membrane stacking of the Golgi, although its precise role in Golgi structure and function is not fully understood yet ^48–52^. We have recently identified GRASP55 as a direct mTOR complex 1 (mTORC1) substrate, establishing the mTORC1-GRASP55 signaling axis as a key metabolic sensory hub that responds to nutrient and stress stimuli to reshape the extracellular proteome accordingly ^7,47,53^.

Here we report that GRASP55, but not GRASP65, is required for proper lysosome function by controlling the trafficking and localization of lysosomal enzymes. Loss of GRASP55 expression leads to missorting and secretion of lysosomal enzymes to the cell culture media, causes lysosome dysfunction and bloating, blunts the lysosomal localization of mTORC1, as well as the phosphorylation of its lysosomal targets TFEB and TFE3, but not that of its non-lysosomal substrates like 4E-BP1. Mechanistically, GRASP55 interacts with GNPTAB and is necessary for its correct ER-Golgi trafficking and protein stability. Consequently, M6P-tagging of lysosomal enzymes is defective in GRASP55 knockout (KO) cells, causing their mislocalization. In sum, these results highlight the role of the Golgi protein sorting machinery in orchestrating Golgi-lysosome communication, lysosome function, and lysosomal mTORC1 signaling. Furthermore, because GRASP55 loss results in a cellular phenotype that is reminiscent of that caused by GNPTAB inactivating mutations, our findings suggest that GRASP55 may be a susceptibility gene in LSD-like conditions.

## Results

### GRASP55 is necessary for proper sorting and delivery of lysosomal enzymes

To gain insight into the role of GRASP55 in the reshaping of the extracellular proteome upon stress, and to reveal the GRASP55-dependent secretome, we previously conducted SILAC-labeling and mass spectrometry (MS)-based proteomics experiments using GRASP55 KO and control WI-26 human lung fibroblasts ^47^ (Fig. 1A). Focusing on the proteins whose secretion is blunted in GRASP55 KO cells, we observed a robust enrichment of extracellular matrix (ECM)- and cell-adhesion-related factors, indicating that GRASP55 controls such cellular processes through the regulation of UPS ^47,53^. Intriguingly, when reanalyzing our secretome data, we noticed the presence of a group of proteins whose secretion is actually enhanced in cells lacking GRASP55 (65 significantly upregulated proteins in the GRASP55 KO cells, compared to the WT controls) (Fig. 1B and Table S1). Gene ontology (GO) term analysis revealed a striking enrichment of lysosomal proteins—among other extracellular proteins—within this dataset (31 out of 65 proteins annotated within the cellular component GO term ‘lysosome’), with most of them being lysosomal enzymes, such as prosaposin (PSAP), cathepsin D (CTSD), cathepsin B (CTSB), β-hexosaminidase (HEXB), and progranulin (PGRN) (Fig. 1B,C and Table S1), in line with a previous study ^54^.

**Figure 1.**
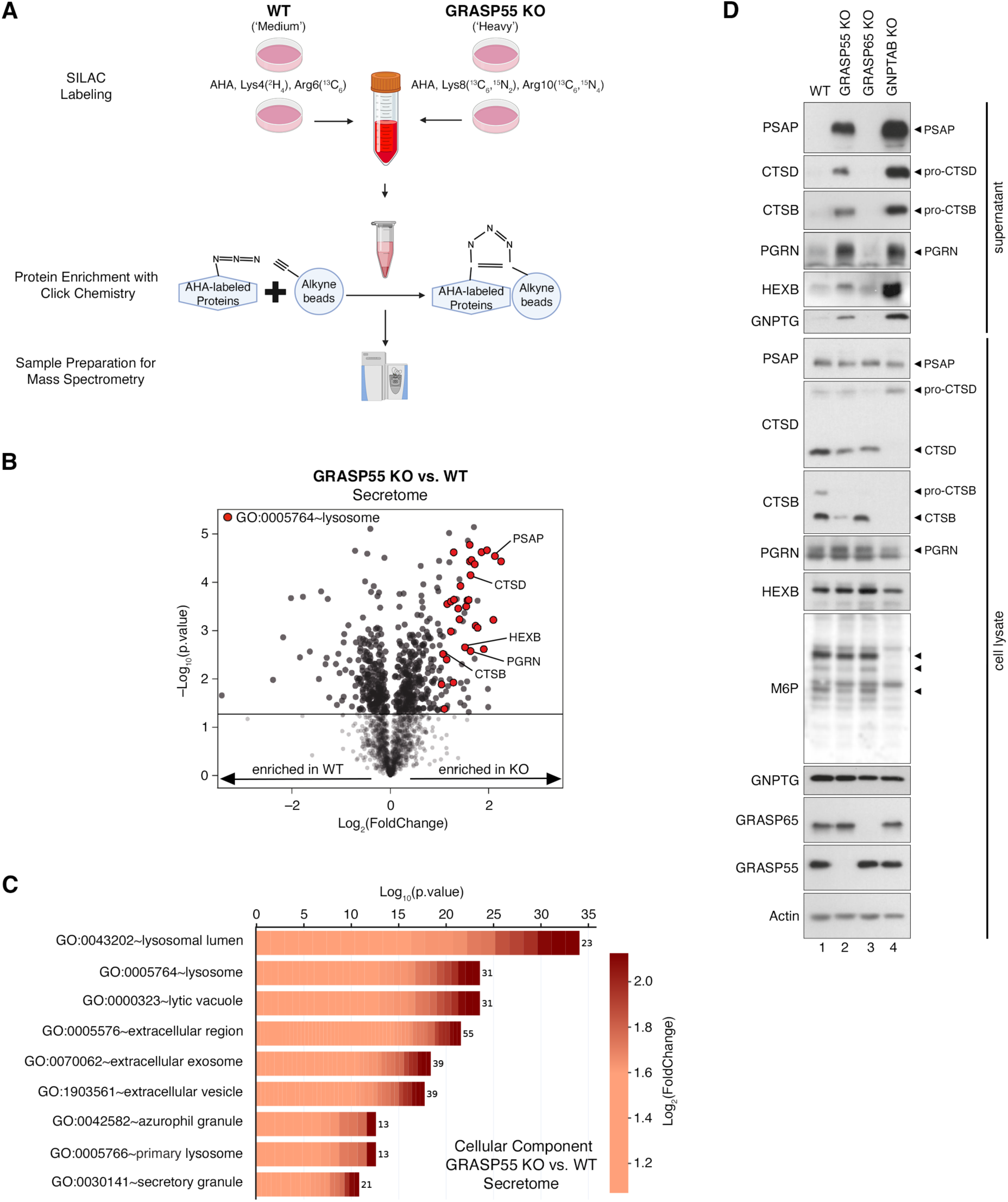
Loss of GRASP55, but not GRASP65, causes missorting of lysosomal enzymes, resembling GNPTAB loss-of-function. **(A)** Experimental outline of the SILAC labeling- and click chemistry-based assay performed to identify the GRASP55-dependent secretome in WI-26 cells (originally described in ^47^; see also Methods). **(B)** Volcano plot showing all proteins identified in the GRASP55-dependent secretome (gray dots). Proteins whose secretion is strongly and significantly increased (Log_2_FC > 1, p<0.05) in GRASP55 KO WI-26 cells were used for the GO analysis shown in (C). Proteins within this subset that belong to the cellular component (CC) GO term ‘lysosome’ are shown in red. **(C)** CC GO term analysis using the proteins whose secretion is enhanced in GRASP55 KO cells reveals strong enrichment of lysosome-related terms. The color of each box represents log_2_-transformed fold change values for each protein in GRASP55 KO vs. WT cells. The number of proteins in the selected dataset for each term is shown on the right side of each bar. **(D)** Immunoblots with the indicated antibodies showing enhanced secretion of lysosomal enzymes in cell culture media of GRASP55 KO and GNPTAB KO, but not GRASP65 KO cells. Secretion of lysosomal proteins (PSAP, CTSD, CTSB, PGRN, HEXB) and GNPTG assayed by immunoblotting in the supernatants of WT, GRASP55 KO, GRASP65 KO, and GNPTAB KO WI-26 cells. Intracellular levels of the same proteins, and of GRASP55, GRASP65, and Actin assayed in whole cell lysates. M6P-modification of intracellular proteins shown as control, with arrowheads indicating bands in the M6P blot that are lost in GNPTAB KO samples. n = 4 independent experiments. See also Figure S1.

Because the aberrant secretion of lysosomal enzymes in GRASP55 KO cells is reminiscent of the phenotypes typically observed in ML-II, we included GNPTAB KO cells in our follow up analyses ^55^. Furthermore, as GRASP65 is structurally very similar to GRASP55, and together they are involved in the maintenance of Golgi morphology ^48^, but is not involved in the regulation of UPS ^47^, we also generated GRASP65 KO cells to test if this abnormal secretory phenotype is specific for GRASP55 or a general characteristic of perturbed Golgi function. In line with our secretome data, immunoblotting analysis of cell culture supernatants and cell lysates from GRASP55 KO and control WI-26 cells showed strongly elevated secretion of the lysosomal enzymes PSAP, CTSD, CTSB, PGRN, and HEXB in the media of GRASP55-deficient cells, similar to that of GNPTAB KO cells (Fig. 1D). Interestingly, in addition to lysosomal enzymes, we also detected GNPTG, the soluble γ subunit of the GNPT complex, in the media of GRASP55 KO cells (and of GNPTAB KO cells) (Fig. 1D). This suggested a potential defect in the assembly and function of the GNPT complex that is necessary for M6P-tagging of lysosomal enzymes ^30,32^. Indeed, the levels of several M6P-modified proteins were greatly reduced in the absence of GRASP55 compared to control cells, albeit not to the extent observed in GNPTAB-deficient cells that completely lack catalytic activity (Fig. 1D). Of note, none of these effects were observed in GRASP65-deficient cells, indicating that GRASP55 controls lysosomal enzyme trafficking and GNPT activity independently of its role in Golgi morphology (Fig. 1D).

In order to investigate the dynamics of lysosomal enzyme sorting and secretion in the absence of GRASP55, we then performed synchronized trafficking experiments using the RUSH (retention using selective hooks) method ^56^, with a streptavidin-KDEL fusion protein as ER luminal hook and PSAP fused to SBP (streptavidin-binding protein). This method allows for the synchronized and rapid release of an SBP-tagged protein from the streptavidin hook by the addition of biotin in the culture medium ^56^. In both WT and GRASP55-deficient cells, PSAP-SBP localized in the ER in the absence of biotin. In WT cells, PSAP-SBP was transported to the Golgi apparatus within 1 hour after biotin addition and was sorted into vesicular structures that did not show substantial overlap with the Golgi at 2-4 hours following biotin addition (Fig. S1A). In contrast, whereas the ER-Golgi trafficking of PSAP-SBP was also observed in GRASP55-deficient cells within 1 hour following biotin addition, the Golgi localization of PSAP-SBP did not change further at later time-points (Fig. S1A), indicating a potential sorting defect in GRASP55 KOs. Therefore, we performed a biotin addition time-course looking at the secretion of PSAP-SBP into the medium of GRASP55 or control cells. As with our experiments assessing the steady-state levels of endogenous PSAP in cell culture supernatants (Fig. 1D), also in RUSH assays PSAP-SBP secretion was strongly elevated in GRASP55-deficient cells, compared to controls (Fig. S1B).

Next, we assessed the role of GRASP55 in the subcellular localization of lysosomal enzymes using confocal microscopy and colocalization analysis of CTSB, CTSD, PSAP, and PGRN with the lysosomal membrane protein LAMP2. Whereas all four proteins colocalized strongly with LAMP2 in control cells, their lysosomal localization was strongly decreased in both GRASP55- and GNPTAB-deficient cells (Fig. 2A-H). Again, this effect was specific for GRASP55, as loss of GRASP65 had no effect in the localization of these enzymes (Fig. 2A-H).

**Figure 2.**
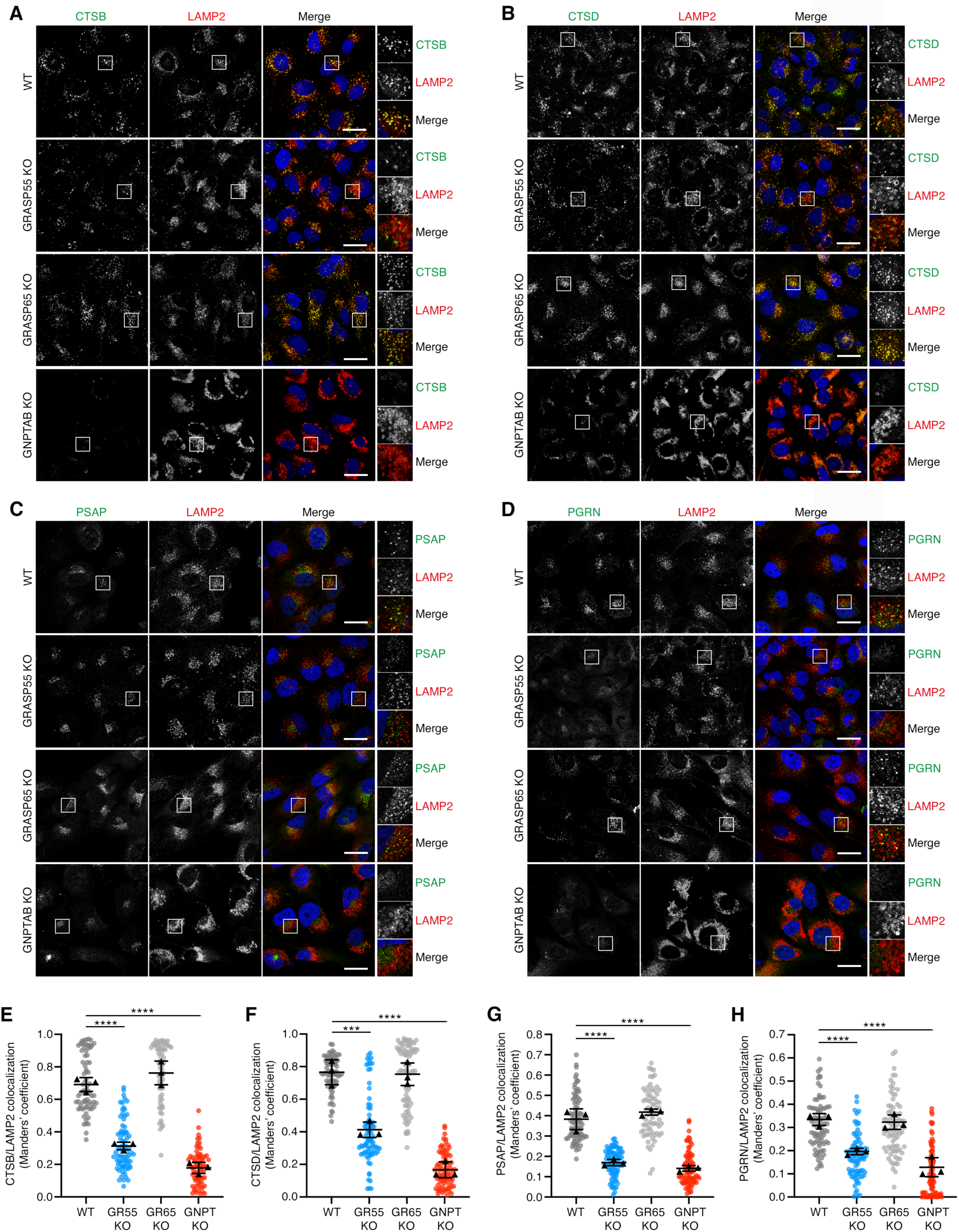
Reduced levels of lysosomal enzymes in lysosomes of GRASP55 KO cells, similar to GNPTAB KO cells. **(A-D)** Colocalization analysis of CTSB (A), CTSD (B), PSAP (C), or PGRN (D) with LAMP2 (lysosomal marker) in WT, GRASP55 KO, GRASP65 KO, and GNPTAB KO WI-26 cells, using confocal microscopy. Nuclei stained with DAPI (blue). Magnified insets shown to the right. Scale bars, 10 μm. **(E-H)** Quantification of colocalization from (A-D). Merged data from 3 independent experiments are shown. n = 70-82 individual cells from 5 independent fields per genotype per experiment. Data in graphs shown as mean ± SD. *** p<0.005, **** p<0.001.

### Cells lacking GRASP55 expression display an LSD-like lysosomal signature

The lysosomal storage disease ML-II, caused by mutations in *GNPTAB*, is typically characterized by the presence of enlarged and defective lysosomes in the diseased cells. More specifically, the inability of lysosomal enzymes to reach their destination leads to the accumulation of non-degraded lysosomal content, which in turn causes an expansion of the lysosomal area and lysosomal bloating. In line with this, we observed diminished lysosomal activity (assayed by Magic Red, a fluorescence-based CTSB activity assay) in GRASP55- and GNPTAB-deficient cells, but not in GRASP65 KO cells (Fig. 3A,B). In addition, using enzymatic activity assays, we detected blunted arylsulfatase B (ARSB), β-hexosaminidase (HEXB), α-L-iduronidase (IDUA), and β-glucuronidase (GUSB) activities in the lysates of GRASP55 and GNPTAB KO cells, accompanied by significantly elevated activities in the respective supernatants (Fig. S2A-H).

**Figure 3.**
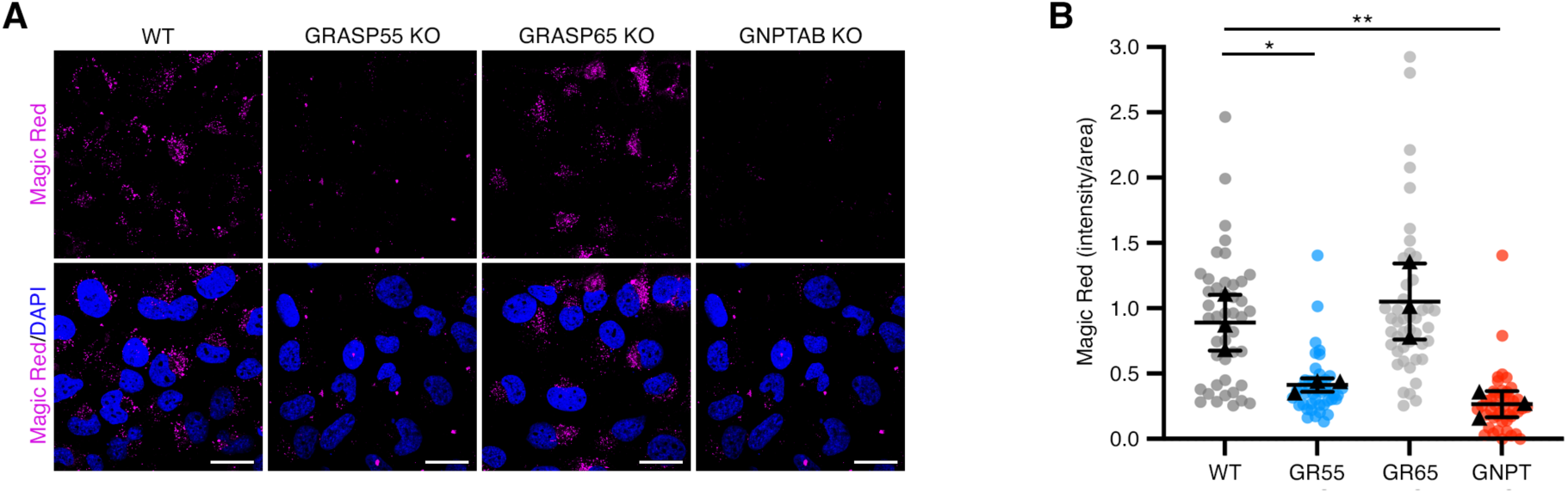
GRASP55-deficient cells contain dysfunctional lysosomes. **(A-B)** Lysosomal cathepsin B (CTSB) activity was monitored using a Magic Red fluorescent assay in WT, GRASP55 KO (GR55 KO), GRASP65 KO (GR65 KO), or GNPTAB KO (GNPT KO) WI-26 cells. The punctate signal is proportional to CTSB proteolytic activity inside lysosomes. Nuclei stained with DAPI. Scale bars, 10 μm (I). Quantification of Magic Red signal (intensity/area) in (J). Merged data from 3 independent experiments are shown. n = 45 cells from 3 independent fields per genotype per experiment. Data shown as mean ± SD. * p < 0.05, ** p < 0.01. See also Figure S2.

Unlike soluble lysosomal enzymes, lysosomal transmembrane proteins do not depend on MPR-dependent pathways for their transportation to lysosomes ^57,58^, therefore we next assessed lysosomal morphology using confocal microscopy and immunofluorescence-based detection of LIMP-2 (lysosomal integral membrane protein-2) and LAMP2 (lysosome-associated membrane protein 2). Indeed, both LIMP-2 and LAMP2 showed a typical punctate lysosomal-like pattern in WT cells, with near-perfect colocalization between the two proteins (Fig. 4A). Further indicating that loss of GRASP55 triggers an LSD-like phenotype, GRASP55-deficient cells contained strongly enlarged lysosomes (Fig. 4A,B), although the effect was less pronounced compared to that observed in GNPTAB KO cells (Fig. 4A,B). Of note, this lysosomal bloating was not observed in GRASP65 KO cells (Fig. 4A,B), underscoring a specific role for GRASP55 in this process.

**Figure 4.**
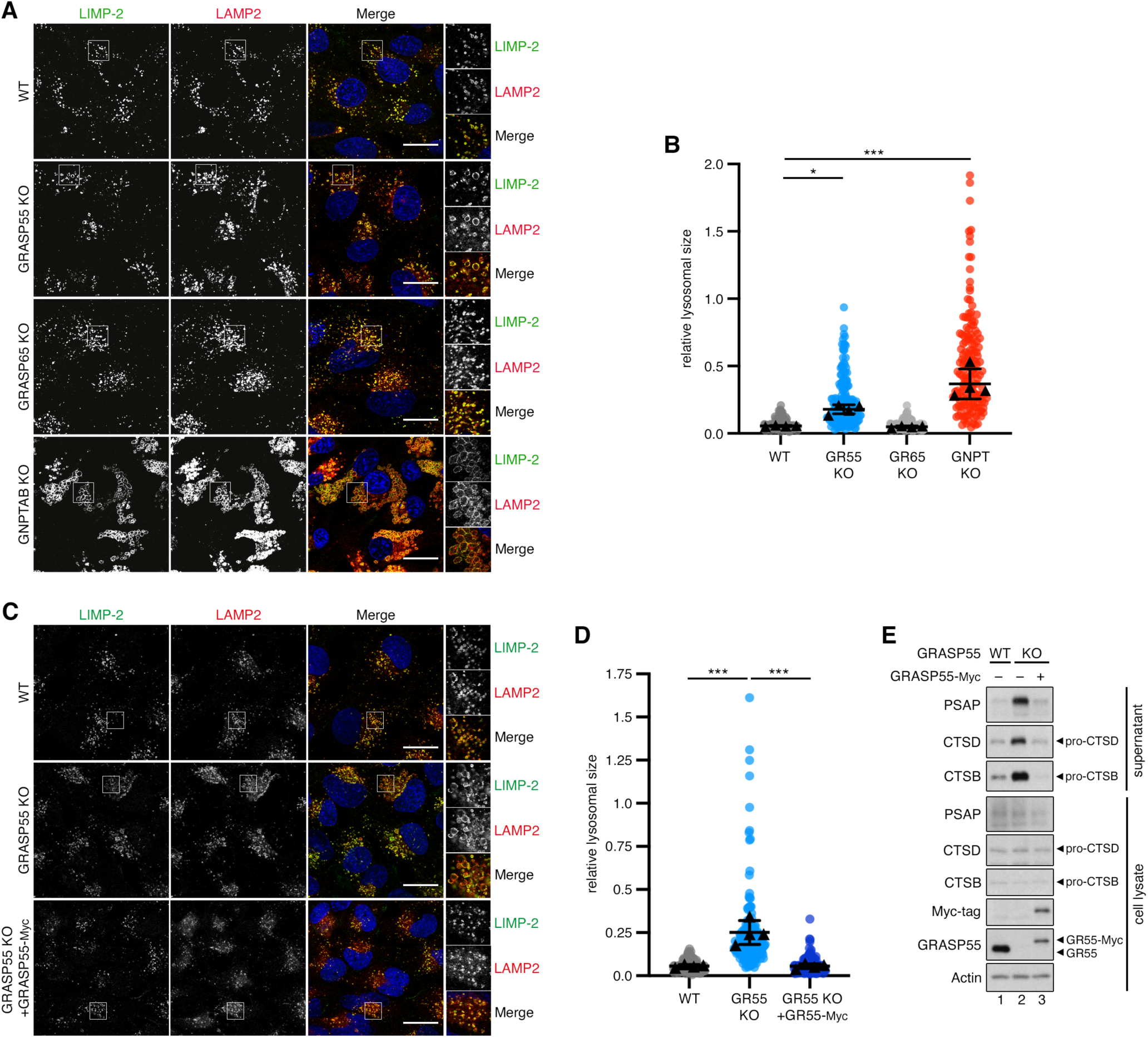
Loss of GRASP55 expression causes lysosomal bloating. **(A-B)** Enlarged lysosome size in GRASP55 KO and GNPTAB KO, but not in GRASP65 KO cells. Immunofluorescence analysis of LIMP-2 and LAMP2 (lysosomal membrane markers) in WT, GRASP55 KO, GRASP65 KO, and GNPTAB KO WI-26 cells, using confocal microscopy. Nuclei stained with DAPI (blue). Magnified insets shown to the right (A). Quantification of relative lysosomal size in (B). Merged data from 4 independent experiments are shown. n = 60-81 individual cells from 5 independent fields per genotype per experiment. **(C-D)** The enlarged lysosome size is reversed by re-expression of GRASP55 in GRASP55 KO cells. Immunofluorescence analysis of LIMP-2 and LAMP2 (lysosomal membrane markers) in WT, GRASP55 KO, and GRASP55 KO WI-26 cells stably expressing Myc-tagged GRASP55, using confocal microscopy. Nuclei stained with DAPI (blue). Magnified insets shown to the right (C). Quantification of relative lysosomal size in (D). Merged data from 4 independent experiments are shown. n = 50-64 individual cells from 5 independent fields per genotype per experiment. **(E)** The aberrant lysosomal enzyme secretion is reversed by re-expression of GRASP55 in GRASP55 KO cells. Secretion of lysosomal enzymes (PSAP, CTSD, CTSB) assayed by immunoblotting in the supernatants of WT, GRASP55 KO, and GRASP55 KO WI-26 cells stably expressing Myc-tagged GRASP55. Intracellular levels of PSAP, CTSD, CTSB, GRASP55, and Actin assayed in whole cell lysates. n = 3 independent experiments. Scale bars, 10 μm. Data in graphs shown as mean ± SD. * p<0.05, *** p<0.005. See also Figure S3.

To ensure that the abnormal lysosomal phenotype is linked to GRASP55 expression, we next reconstituted the GRASP55 KO WI-26 cells by stably re-expressing Myc-tagged GRASP55. Exogenously expressed GRASP55 localized properly at the Golgi, similarly to endogenous GRASP55 localization in WT cells (Fig. S3). Importantly, reconstitution of GRASP55 expression fully reversed the lysosomal bloating (Fig. 4C,D) and the lysosomal enzyme secretion phenotypes (Fig. 4E) observed in GRASP55 KO cells. In sum, these data indicate the specific role of GRASP55 in the sorting and trafficking of lysosomal enzymes and in the maintenance of proper lysosome function, while GRASP55 deletion resembles the LSD-like phenotypes that are driven by loss of GNPTAB expression.

### Lysosomal dysfunction upon loss of GRASP55 selectively perturbs lysosomal mTORC1 signaling

Active mTORC1 controls virtually all cellular functions through the phosphorylation of multiple effector proteins that takes place at various subcellular locations ^14,16^. For instance, while mTORC1 regulates protein synthesis by phosphorylating S6K and 4E-BP1 in the cytoplasm, it phosphorylates TFEB and TFE3 on the lysosomal surface to prevent their nuclear translocation and block lysosome biogenesis ^7,59^. Furthermore, we recently showed that pharmacological or genetic blockage of basal lysosomal proteolysis (e.g., by BafA1 treatment; or in GNPTAB KO cells) specifically affects the phosphorylation of lysosomal mTORC1 substrates, as it causes its relocalization away from the lysosomal surface without downregulating its activity towards its non-lysosomal substrates ^7^. Therefore, we next assessed the localization and activity of mTORC1 in GRASP55 KO cells. As observed previously in GNPTAB KO HEK293FT cells ^7^, the lysosomal localization of mTOR was diminished in GRASP55 KO and GNPTAB KO, but not in GRASP65 KO cells, resembling the pattern seen upon treatment with amino acid (AA) starvation media (Fig. 5A,B). Accordingly, loss of GRASP55 or GNPTAB expression blunted the phosphorylation of the lysosomal mTORC1 substrates, TFEB and TFE3, but not that of the cytoplasmic substrate 4E-BP1 (Fig. 5C). As a control, treatment with Torin1, an ATP-competitive mTOR inhibitor, caused diminished phosphorylation of all mTORC1 substrates (Fig. 5C). Consistent with the changes in its phosphorylation, we observed enhanced nuclear translocation of TFE3 in GRASP55 KO and GNPTAB KO cells (as in all Torin1-treated cells), but not in GRASP65 KOs (Fig. 5E,F). Taken together, these data show that the defect in lysosomal function that is observed in GRASP55- and GNPTAB-deficient cells selectively perturbs lysosomal mTORC1 signaling, without affecting its activity towards its cytoplasmic targets, like 4E-BP1.

**Figure 5.**
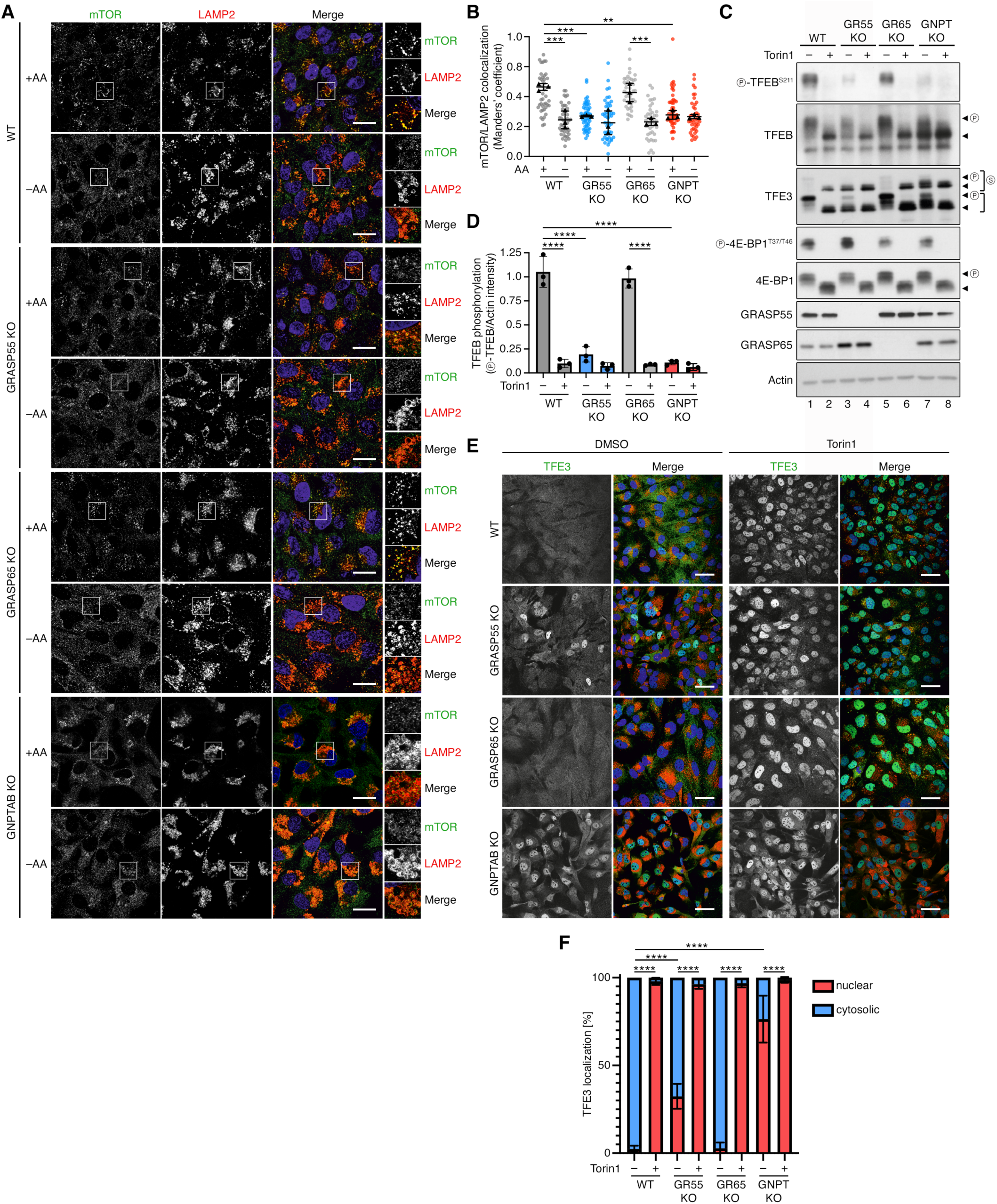
Lysosomal dysfunction in GRASP55 and GNPTAB KO cells specifically perturbs lysosomal mTORC1 signaling. **(A-B)** Loss of lysosomal mTOR localization in GRASP55 KO and GNPTAB KO cells. Colocalization analysis of mTOR with LAMP2 (lysosomal marker) in WT, GRASP55 KO, GRASP65 KO, and GNPTAB KO WI-26 cells, treated with media containing (+AA) or lacking AAs (–AA) for 2 hours, using confocal microscopy. Nuclei stained with DAPI (blue). Magnified insets shown to the right. Scale bars, 10 μm (A). Quantification of colocalization in (B). Merged data from 3 independent experiments are shown. n = 60 individual cells from 5 independent fields per genotype per experiment. **(C-D)** Diminished phosphorylation of TFEB/TFE3, but not 4E-BP1, in GRASP55 KO or GNPTAB KO cells. Immunoblots with lysates from WT, GRASP55 KO, GRASP65 KO, and GNPTAB KO WI-26 cells, treated with DMSO (vehicle) or Torin1 (250 nM) for 2 hours, and probed with the indicated antibodies (C). Quantification of TFEB phosphorylation (p-TFEB/Actin ratio) in (D). Arrowheads indicate bands corresponding to different protein forms, when multiple bands are present. P: phosphorylated form, S: SUMOylated form. n = 3 independent experiments. **(E-F)** Nuclear translocation of TFE3 in GRASP55 KO or GNPTAB KO cells. TFE3 localization analysis in WT, GRASP55 KO, GRASP65 KO, and GNPTAB KO WI-26 cells, treated with DMSO (vehicle) or Torin1 (250 nM) for 2 hours, using confocal microscopy. LAMP2 used as lysosomal marker (red). Nuclei stained with DAPI (blue). Magnified insets shown to the right. Scale bars, 20 μm (E). Quantification of % nuclear/cytosolic TFE3 localization in (F). Merged data from 4 independent experiments are shown. n = 80-118 individual cells from 5 independent fields per genotype per experiment. Data in graphs shown as mean ± SD. ** p<0.01, *** p<0.005, **** p<0.001.

### GRASP55 controls GNPTAB localization and protein stability

We next aimed to investigate the underlying cause for the missorting of lysosomal enzymes in GRASP55-deficient cells. As we observed an overall reduction of M6P protein modification, as well as abnormal secretion of GNPTG, in GRASP55 KO cells (Fig. 1D), we focused our analysis on GNPTAB, the catalytic part of the GNPT complex. As expected, exogenously expressed, Myc-tagged GNPTAB showed a near-perfect colocalization with the *cis*-Golgi marker GM130 ^60^ in control cells (Fig. 6A). Interestingly, however, loss of GRASP55 led to strongly reduced Golgi localization of GNPTAB (Fig. 6A,B), accompanied by increased colocalization with PDI (Protein disulfide-isomerase), a marker of the ER lumen ^61^ (Fig. 6C,D), indicating that GRASP55 is necessary for maintaining GNPTAB at the Golgi apparatus. Indeed, in co-immunoprecipitation (co-IP) experiments, we detected a reciprocal interaction between GNPTAB and endogenous GRASP55, but not with the closely-related GRASP65 or its interactor GM130 (Fig. 6E). Furthermore, we did not detect binding between GNPTAB and the GRASP55 binding partner, GOLGIN-45, suggesting that the GNPTAB-GRASP55 interaction is direct and specific (Fig. 6E).

**Figure 6.**
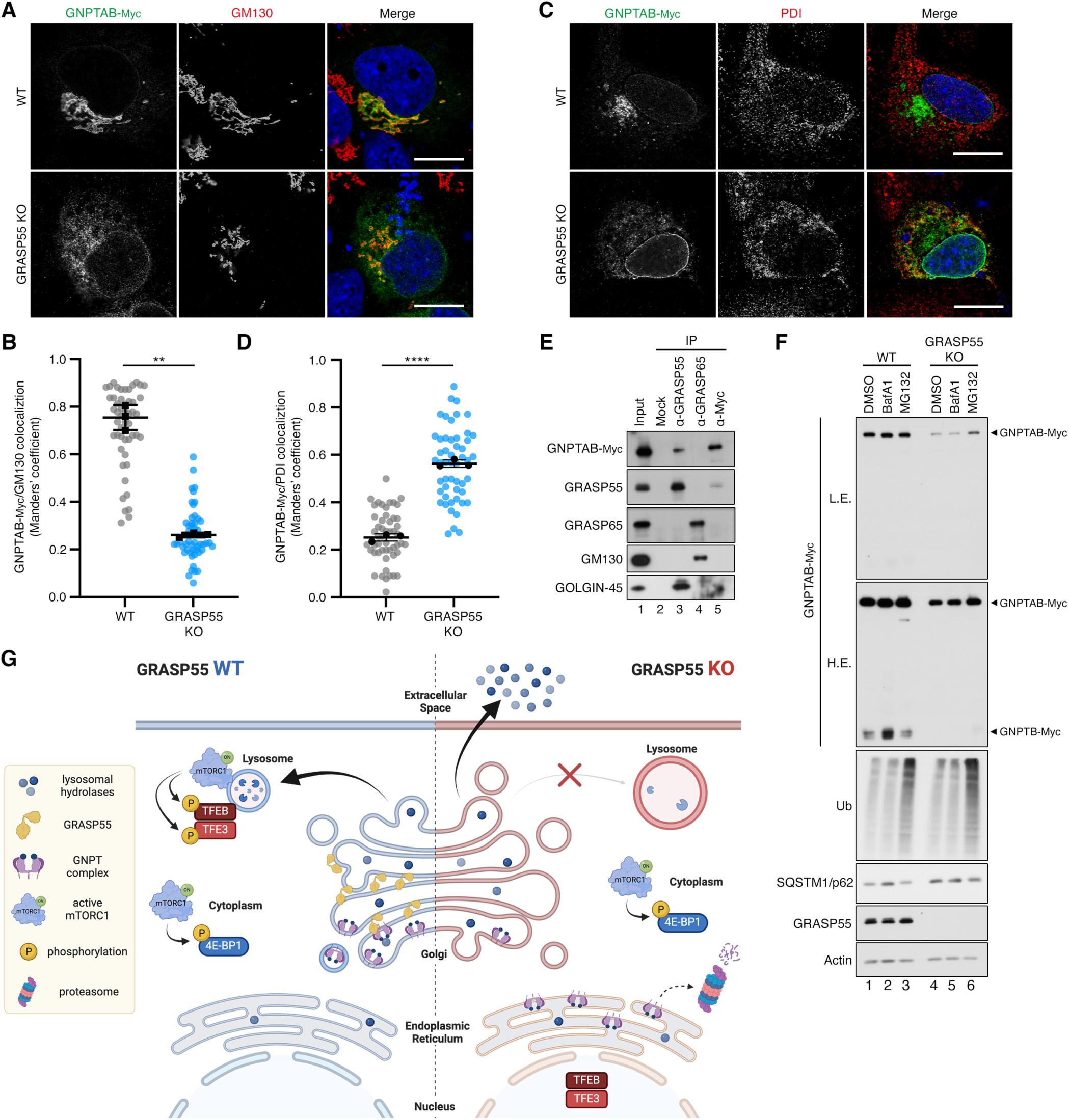
GRASP55 is required for proper GNPTAB localization at the Golgi and protein stability. **(A-B)** Blunted Golgi localization of GNPTAB in GRASP55 KO cells. Colocalization analysis of exogenously expressed Myc-tagged GNPTAB with GM130 (Golgi marker) in WT or GRASP55 KO WI-26 cells, using confocal microscopy. Nuclei stained with DAPI (blue). Scale bars, 10 μm (A). Quantification of colocalization in (B). Merged data from 3 independent experiments are shown. n = 50 individual cells from 5 independent fields per genotype per experiment. Data shown as mean ± SD. ** p<0.01. **(C-D)** Aberrant ER localization of GNPTAB in GRASP55 KO cells. Experiment as in (A-B), but for GNPTAB colocalization with PDI (ER marker) (C). Quantification of colocalization in (D). Merged data from 3 independent experiments are shown. n = 51 individual cells from 5 independent fields per genotype per experiment. Data shown as mean ± SD. **** p<0.001. **(E)** GRASP55, but not GRASP65, interacts with GNPTAB. Co-immunoprecipitation analysis of endogenous GRASP55, GRASP65, and exogenously expressed Myc-tagged GNPTAB proteins in WI-26 cells. The input and IP samples were analyzed by immunoblotting using antibodies against the indicated proteins. The GRASP55 interactor GM130 and the GRASP65 interactor GOLGIN-45 were used as positive controls. n = 3 independent experiments. **(F)** Decreased stability of GNPTAB is partially rescued by proteasomal inhibition. Immunoblots with lysates from WT or GRASP55 KO WI-26 cells, transiently expressing Myc-tagged GNPTAB, treated with DMSO (vehicle), BafA1 (100 nM, 8h), or MG132 (10 μM, 8h), and probed with the indicated antibodies. L.E.: low exposure; H.E.: high exposure. n = 3 independent experiments. **(G)** Proposed model for the role of GRASP55 in proper sorting and of lysosomal enzymes at the Golgi via the regulation of ER-Golgi trafficking of GNPTAB. As a result, lysosomal enzymes are aberrantly secreted in GRASP55 KO cells, which exhibit dysfunctional and bloated lysosomes, resembling LSD-like phenotypes. The lysosomal dysfunction in GARSP55 KO cells specifically perturbs lysosomal mTORC1 signaling, without affecting cytoplasmic mTORC1 activity, causing nuclear translocation and activation of the TFEB/TFE3 transcription factors.

At the *cis*-Golgi, the GNPTAB precursor protein undergoes a maturation process, activated through proteolytic cleavage by S1P to the enzymatically active GNPTA and GNPTB subunits ^31^. Indeed, in control WI-26 cells, a fraction of the exogenously expressed GNPTAB was processed, as shown by the presence of the cleaved Myc-tagged GNPTB (Fig. 6F). In contrast, consistent with the mislocalization and reduced Golgi presence of GNPTAB in GRASP55 KO cells, we also observed defective GNPTAB processing (Fig. 6F, lanes 1 and 4). Strikingly, these experiments also revealed reduced expression of Myc-tagged GNPTAB in GRASP55 KO cells (Fig. 6F, lanes 1 and 4), which was partially rescued by treatment with the proteasome inhibitor MG132, but not with the lysosome inhibitor BafA1 (Fig. 6F, lanes 5-7). In sum, these results indicate that GRASP55 is necessary for proper retention and processing of GNPTAB at the Golgi, presumably via direct interaction between the two proteins. Moreover, in GRASP55-deficient cells, GNPTAB is mislocalized to the ER where it is targeted for degradation, in part through proteasome-mediated mechanisms (Fig. 6G).

## Discussion

### GRASP55 as a master regulator of protein sorting at the Golgi

Initial work on the closely-related GRASP55 and GRASP65 proteins proposed their involvement in the stacking of Golgi cisternae, however their precise role in Golgi assembly and morphology has been debated by other studies ^45,48–52^. Whereas GRASP55 and GRASP65 share high amino acid sequence similarity in their N-terminal halves, the respective C-terminal serine/proline-rich (SPR) regions are substantially different between the two proteins ^45,50^, suggesting that they are also involved in distinct cellular functions. Indeed, we have recently demonstrated that GRASP55, independently of GRASP65, regulates UPS downstream of mTORC1 and starvation/stress signaling ^46,47,53^; and that it controls the intra-Golgi distribution and the compartmentalized localization of glycosylation enzymes that are responsible for sphingolipid biosynthesis ^62^. In this study, by reanalyzing our previously published GRASP55-dependent secretome dataset ^47^, we reveal a newfound important role for GRASP55 in proper sorting of lysosomal enzymes at the Golgi, with GRASP55-deficient cells showing a typical LSD-like phenotype, manifested as lysosomal enzyme missorting and secretion, lysosomal bloating, and impaired lysosome function. Of note, our data highlight a specific role for GRASP55 in this process, as GRASP65 loss-of-function cells show normal lysosomal enzyme sorting and lysosomal function.

### GRASP55 lies at the center of mTORC1 signaling

We previously found that mTORC1 directly phosphorylates a fraction of GRASP55 at the Golgi to control its subcellular localization and prevent the activation of UPS ^47,53^. Accordingly, in TSC-deficient cells, a well-described model of mTORC1 hyperactivation, GRASP55 remained phosphorylated and did not relocalize away from the Golgi even in starved or stressed cells ^47,53^. Besides its role in UPS, hyperactivation of mTORC1 in cellular models of LSDs (assayed by phosphorylation of its canonical substrate S6K) was suggested to be involved in the pathogenesis of the disease due to suppression of autophagy ^63^. In addition, mTOR signaling may also play a role in LSDs by controlling lysosomal biogenesis, via the phosphorylation of its direct targets, the TFEB/TFE3 transcription factors ^22,64–67^. In particular, TFEB and TFE3 regulate the expression of lysosomal enzymes, lysosomal membrane proteins, as well as lysosome protein receptors ^22^. Interestingly, a recent study showed that TSC-mutant cells have more and enlarged lysosomes due to aberrant activation of TFEB/TFE3 transcription factors ^68,69^, raising the plausible hypothesis that this TSC-mTORC1-TFEB/TFE3 pathway may be regulating lysosomal biogenesis partly also via the hyperphosphorylation of GRASP55. Whether the GRASP55-mediated lysosomal protein sorting is also actively regulated downstream of nutrient and stress signaling will require additional work and remains to be seen in future studies.

A major site of mTORC1 activation by nutrients is the lysosomal surface, where it is recruited by the Rag GTPase heterodimers when nutrients are abundant and lysosomes are functional ^5–7^. While the lysosomal localization of mTORC1 is absolutely necessary for the phosphorylation of its lysosomal substrates like TFEB and TFE3, we recently showed that, in the presence of exogenous amino acids, non-lysosomal mTORC1 is also active and can potently phosphorylate its cytoplasmic (e.g., S6K, 4E-BP1) or Golgi-localized substrates like GRASP55 ^7^. Furthermore, perturbing the sorting and delivery of lysosomal enzymes by GNPTAB knockout or knockdown caused delocalization of mTORC1 from the lysosomal surface and specifically affected the phosphorylation of its lysosomal substrates, indicating that lysosome dysfunction differentially affects substrate specificity downstream of mTORC1 ^7^. Notably, similar effects were observed in this study using GRASP55-deficient cells, which exhibited diminished lysosomal mTOR localization and enhanced dephosphorylation and activation of the TFEB/TFE3 transcription factors. This aberrant activation of lysosomal biogenesis pathways is a particularly interesting—but poorly understood—aspect of LSDs, as it seemingly triggers a vicious circle in cells with dysfunctional lysosomes which promote the generation of even more dysfunctional lysosomes. This is readily evident in classical cellular models of LSDs (e.g., GNPTAB-mutant cells) or in GRASP55 KO cells, in which more and larger lysosomes occupy a large part of the cell volume. Whether blocking this feed-forward loop (e.g., by reducing the expression or blocking the activation of TFEB/TFE3) is beneficial for cell growth and for the homeostatic response of cells to nutrient starvation, two parameters that strongly rely on proper lysosomal function, is currently not clear and warrants further investigation in follow-up studies.

In sum, distinct Golgi-based mechanisms play a central role in the activation of mTORC1 by nutrients on the lysosomal surface, thereby fine-tuning metabolic signaling and organelle biogenesis in cells. In turn, mTORC1 activity may regulate Golgi function and protein secretion via the phosphorylation of key targets like GRASP55. Such feedback loops are commonly found in the mTORC1 signaling network. Characteristic examples are the regulation of protein synthesis via the mTORC1-dependent phosphorylation of core components of the translation initiation machinery, with eIF4a/EIF4A functioning also as a negative upstream regulator of mTORC1 in *Drosophila* and mammalian cells ^70^; or the direct inhibition of mTORC1 by malonyl-CoA, a metabolic intermediate in de novo lipid biosynthesis that is produced by ACC1 (Acetyl-CoA carboxylase) and is consumed by FASN (fatty acid synthase), two enzymes whose expression levels are, in turn, regulated downstream of mTORC1 and the SREBP transcription factors ^71,72^.

### GRASP55 is required for proper trafficking of GNPTAB

The heterohexameric GNPT complex localizes at the *cis*-Golgi and is responsible for the M6P-tagging of lysosomal enzymes, enabling their proper sorting at the *trans*-Golgi network (TGN) and their trafficking to the late endosomal compartment. Loss of GNPT expression, activity, or defective processing of the GNPTAB precursor causes ML-II in humans. Here we show that GRASP55, but not GRASP65, interacts with non-processed GNPTAB and is necessary for its proper localization at the *cis*-Golgi, and for efficient M6P-tagging of the lysosomal enzymes. In the absence of GRASP55 expression, GNPTAB is mislocalized at the ER, a phenomenon that is accompanied by reduced GNPTAB processing, reduced complex stability, and enhanced secretion of the soluble GNPTG subunit. Of note, similar effects on GNPTAB processing and stability were observed in cells lacking expression of LYSET/TMEM251, another Golgi-based factor that was recently shown to be necessary for proper trafficking of lysosomal enzymes ^73–77^.

Taken together, our data indicate that GRASP55 controls the sorting of lysosomal enzymes by ensuring the correct ER-to-Golgi and intra-Golgi trafficking of GNPTAB. Intriguingly, our recently published GRASP55 proximome analysis ^47^ indicated that GRASP55 may be interacting with specific COG (conserved oligomeric Golgi) complex proteins, which play important roles in vesicle and cargo trafficking at the Golgi. Furthermore, mutations in most COG complex subunits are linked to congenital disorders of glycosylation (COG-CDG) and COG-deficient cells are characterized also by the presence of enlarged endolysosomal structures ^78,79^. Future work will be necessary to investigate if and how the interaction of GRASP55 with COG proteins and GNPTAB may be influencing the sorting and secretion of lysosomal proteins.

### Highlighting the Golgi-Lysosome axis in lysosome biology and in LSDs

Although the role of the Golgi apparatus as the main cargo distribution center in cells is well recognized, the underlying molecular mechanisms of cargo selection remain very poorly understood. With regards to lysosomal enzyme sorting, only a handful of proteins, like GNPTAB, the MPRs, or LYSET, were found to participate in the tagging and trafficking of these enzymes, while even fewer have been linked to human LSD-like syndromes when mutated in patients (in addition to mutations in the lysosomal enzymes themselves) ^41,42,77^. Here, we reveal a tight connection between GRASP55 and lysosomal enzyme sorting at the Golgi, underscore the importance of this Golgi-lysosome communication for health and disease, and highlight *GRASP55/GORASP2* as a putative susceptibility gene in LSD-like diseases. The central role of GRASP55 in the regulation of multiple key cellular processes at the Golgi and elsewhere, as well as its extensive protein interaction network, suggest that additional, yet unidentified proteins and molecular mechanisms at the Golgi are likely relevant for the maintenance of lysosomal function and the fine-tuning of metabolic signaling in cells.

In sum, a better characterization of the molecular interplay between nutrient sensing signaling pathways, lysosomal protein sorting at the Golgi, and lysosomal function will be important to understand how these machineries coordinate to control multiple physiological processes in cells, and what goes wrong in associated diseases in humans.

## Methods

### Cell culture and treatments

All cell lines were grown at 37°C, 5% CO_2_. Human male diploid lung WI-26 SV40 fibroblasts (WI-26 cells; #CCL-95.1, ATCC; RRID: CVCL_2758) were cultured in DMEM/F12 GlutaMAX medium (#31331093, Thermo Fisher Scientific), containing 10% FBS (FBS.HP.0500, Bio&SELL) and 1% Pen/Strep (P4333-100ML, Sigma-Aldrich). The identity of the WI-26 cells was validated using the Short Tandem Repeat (STR) profiling service, provided by Multiplexion GmbH. No commonly misidentified cell lines were used in this study. All cell lines were regularly tested for *Mycoplasma* contamination, using a PCR-based approach and were confirmed to be *Mycoplasma*-free.

For amino acid (AA) starvation experiments, culture media were replaced with AA-free DMEM/F12 (D9811-01, US Biologicals) containing 10% dialyzed FBS (F0392, Sigma-Aldrich) for 2 h. To inhibit mTOR kinase activity, cells were treated with 250 nM Torin1 (Cay10997-10, Cayman Chemical) for 2 hours. For Bafilomycin A1 (#BML-CM110-0100, Enzo Life Sciences) treatment, the drug was added to a final concentration of 100 nM in the media for 8 hours before lysis. For proteasome inhibition, cells were treated with 20 μM MG132 (S2163, Selleckchem) for 8 h before lysis. For all drug treatments, DMSO (#4720.1, Roth) was used as vehicle control. Treatments were performed by replacing the culture media with drug-containing media.

### Antibodies

A list of all primary antibodies used in this study is found in Table S2. The H4B4 antibody against LAMP2 was obtained from the Developmental Studies Hybridoma Bank, created by the NICHD of the NIH and maintained at The University of Iowa, Department of Biology. H4B4 was deposited to the DSHB by August, J.T. / Hildreth, J.E.K. (DSHB Hybridoma Product H4B4) ^80^.

### Plasmids and molecular cloning

Plasmid expression vectors were generated by cloning PCR-amplified cDNAs, using appropriate primers. All primer sequences are listed in Table S3. For cDNA generation, total RNA was isolated from human WI-26 fibroblasts using a standard Trizol/chloroform-based extraction (15596018, Thermo Fisher Scientific), and converted to cDNA using Superscript II (#18064014, Thermo Fisher Scientific). For the construction of the GNPTAB expression vector pcDNA4/TO/hGNPTAB-Myc-His, human GNPTAB (NM_024312.5) was amplified from WI-26 cDNA and cloned into the BamHI/NotI restriction sites of the pcDNA4/TO/Myc-His-A plasmid (V103020, Invitrogen). The RUSH construct for PSAP (pIRES-Str-KDEL-hPSAP-SBP) was generated by amplification of human PSAP (NM_002778.4) from WI-26 cDNA and cloning of the cDNA into the AscI and XhoI restriction sites of the Str-KDEL_ManII-SBP-EGFP vector (Addgene plasmid #65252, described in ^56^). Next, the SBP (Streptavidin-Binding Peptide) tag was amplified from Str-KDEL_ManII-SBP-EGFP and inserted in-frame with PSAP using the XhoI and PacI restriction sites.

The pSpCas9(BB)-2A-Puro (PX459) V2.0 plasmid (Addgene plasmid #62988, described in ^81^) was used for the generation of GRASP65/GORASP1- and GNPTAB-deficient WI-26 cells. In brief, double-stranded DNA oligos that encode guide RNAs (gRNAs) against target genes were cloned into the BbsI restriction sites of the PX459 vector. The oligo sequences used for the sgRNA expression plasmids to generate the GRASP65 and GNPTAB KO lines are provided in Table S3.

All restriction enzymes were purchased from New England Biolabs. The integrity of all constructs was verified by sequencing.

### Plasmid DNA transfections

Plasmid DNA transfections were performed using the X-tremeGENE HP DNA transfection reagent (#06366236001, Roche) in a 2:1 DNA/transfection reagent ratio when the cells reached approximately 70% confluency, according to the manufacturer’s instructions. Twenty-four hours post-transfection, cells were either lysed for immunoblotting or fixed for immunofluorescence.

### Generation of knock-out cell lines

The GRASP55 KO WI-26 cells were described previously ^47^. The GRASP65 and GNPTAB knockout cell lines were generated using the PX459-based CRISPR/Cas9 method, as described elsewhere ^81^. In brief, cells were transfected with sgRNA-expressing vectors and 36 hours post-transfection they were selected with puromycin (2 μg/ml) (#A1113803, Gibco) for 5 days. Single cell clones were picked using cloning cylinders (#CLS31668, Sigma-Aldrich) and knockout clones were validated by genomic DNA sequencing and immunoblotting. An empty PX459 vector was used to generate matching control cell lines.

### Generation of stable cell lines

A reconstituted GRASP55 KO WI-26 cell line, stably expressing Myc-tagged human GRASP55 was described previously ^7^.

### Cell lysis, preparation of supernatant samples, and Immunoblotting

For standard SDS-PAGE and immunoblotting experiments, cells from one well of a 6-well plate, at approximately 90% confluence, were lysed in-well with 300 μl ice-cold Triton lysis buffer (50 mM Tris-HCl pH 7.5, 0.5% Triton X-100, 150 mM NaCl, 0.1% SDS), supplemented with 1x EDTA-free cOmplete protease inhibitors (#11873580001, Roche) and 1x PhosSTOP phosphatase inhibitors (#4906837001, Roche). Lysates were clarified by centrifugation (12,000 x g, 15 min, 4°C) to remove debris and supernatants transferred to a new tube. Samples were boiled in 1x SDS sample buffer for 5 min at 95 °C (6x SDS sample buffer: 350 mM Tris-HCl pH 6.8, 30% glycerol, 600 mM DTT, 12.8% SDS, 0.12% bromophenol blue).

For protein secretion experiments, serum-free supernatants were centrifuged (2000 x g, 5 min, 4°C) to remove dead cells and debris. Cleared culture supernatants were concentrated using 3 kDa cut-off concentrator tubes (#516-0227P, VWR) according to the manufacturer’s instructions, and Laemmli loading buffer (1x final concentration; 4x Laemmli sample buffer composition: 250 mM Tris-HCl pH 6.8, 8% SDS, 40% glycerol, 200 mM DTT, 0.02% bromophenol blue) was added to the concentrated supernatants.

Protein samples were subjected to electrophoretic separation on SDS-PAGE and analysed by standard Western blotting techniques. In brief, proteins were transferred to PVDF membranes (IPVH00010, Millipore), except for M6P and phospho-TFEB^S211^ blots for which nitrocellulose membranes (#10600002, Amersham) were used instead. Membranes were blocked with 5% skim milk powder (#42590, Serva) in TBS-T buffer [50 mM Tris-HCl pH 7.4, 150 mM NaCl, 0.1% Tween-20 (#A1389, AppliChem)] or with 5% BSA (#10735086001, Roche; #8076, Carl Roth) in TBS-T for phosphoprotein immunoblots, and incubated with primary antibodies diluted in TBS-T, for 1 hour at room temperature or overnight at 4°C, followed by incubation with appropriate HRP-conjugated secondary antibodies for 1 hour at room temperature. For the phospho-TFEB blots, the primary antibody was incubated in TBS-T also containing 5 % BSA. Signals were detected by enhanced chemiluminescence (ECL) and immunoblot images were captured on films (#28906835, GE Healthcare; #4741019289, Fujifilm). Quantification of immunoblots was performed by densitometric analysis of the band intensities, using the Gel analysis tool of the ImageJ software ^82^. A list of all primary and secondary antibodies used in this study is provided in Table S2.

### Co-immunoprecipitation (co-IP)

For co-IP experiments, of WI-26 cells (1x 10 cm dish per condition) were transfected with a GNPTAB-Myc expression vector. Cells were lysed 24 hours post-transfection in 500 μl Triton lysis buffer per dish (50 mM Tris-HCl pH 7.5, 1% Triton X-100, 150 mM NaCl, 1x EDTA-free cOmplete protease inhibitors, 1x PhosSTOP phosphatase inhibitors) using a glass Dounce homogenizer and incubated for 30 minutes on ice. Samples were clarified by centrifugation (12,000 x g, 10 min, 4°C) and incubated with 2 μg of anti-GRASP55 monoclonal antibody, 2 μg of anti-GRASP65, or 5 μl of mouse monoclonal anti-Myc-tag antibody for 3-4 hours at 4 °C under constant agitation. Complexes were incubated overnight with 40 μl protein G agarose beads, washed 4x with lysis buffer, boiled in 1x Laemmli sample buffer (5 min, 95 °C), and separated by SDS-PAGE. Input samples (50 μl) were collected before pre-clearing, Laemmli loading buffer (1x final concentration) was added to the lysates, samples were boiled and analyzed by immunoblotting.

### Immunofluorescence and confocal microscopy

Immunofluorescence/confocal microscopy experiments were performed as described previously ^9,11,47^ with minor modifications. In brief, cells were grown on glass coverslips and treated as described in the figure legends. Samples were fixed for 10 min at room temperature with 4% formaldehyde (#252549, Sigma-Aldrich) in PBS (for the experiments shown in figures 2C, 2D, 3, 6, S2 and S3) or for 5 min at –20 °C with 100% ice-cold methanol (for the experiments shown in figures 2A, 2B, 4, 5), permeabilized with 0.5% NP-40 (I8896, Sigma-Aldrich) in PBS for 10 min, blocked with 1% FBS (FBS.HP.0500, Bio&SELL) in PBS for 30 min, incubated with primary antibodies diluted in blocking buffer (1% FBS in PBS) for 1 hour at room temperature, washed 3x with blocking buffer, and incubated with appropriate highly cross-adsorbed secondary antibodies conjugated to Alexa Fluor 488 or 555 (Thermo Fisher Scientific) for 1 hour at room temperature. Nuclei were stained with 0.1 μg/ml DAPI (#D9542, Sigma-Aldrich). Samples were mounted with Fluoromount-G mounting medium (#00-4958-02, Invitrogen). All images were acquired on an SP8 Leica confocal microscope (TCS SP8 X or TCS SP8 DLS, Leica Microsystems) using 63x or 100x objective lenses. Image acquisition was performed using the LAS X software (Leica Microsystems). Images from single channels are shown in grayscale, whereas in merged images, Alexa Fluor 488 is shown in green and Alexa Fluor 555 in red. Brightness and contrast were adjusted for visualization purposes using Fiji (https://imagej.net/software/fiji/downloads) ^83^. Alterations were applied to the entire image, keeping the parameters identical between all images of the same channel in each panel.

### Quantification of colocalization

Colocalization analysis in confocal microscopy experiments was performed as in ^11,13^, using the Coloc2 plugin of the Fiji software ^83^. At least 50 individual cells from 5 independent representative images per condition per replicate (3-4 replicates per experiment) were used to calculate Manders’ colocalization coefficient (MCC) with automatic Costes thresholding ^84–86^. Outlines of individual cells were traced, excluding the area corresponding to the cell nucleus, to generate the region of interest (ROI) used for calculating the MCC to prevent false-positive colocalization due to automatic signal adjustments. MCC is defined as a part of the signal of interest (lysosomal enzymes, mTOR, or GNPTAB-myc), which overlaps with a second signal (LAMP2).

### Enzymatic activity measurements

The enzymatic activities of the lysosomal enzymes arylsulfatase B (ARSB), β-hexosaminidase (HEXB), α-iduronidase (IDUA) and β-glucuronidase (GUSB) in cell lysates and corresponding serum-free supernatants of cultured cells were assayed by estimation of 4-nitrophenol or 4-methylumbelliferone liberated from the enzyme-specific substrate ^55,87,88^.

### Magic Red lysosomal protease activity assay

Intracellular lysosomal protease activity was measured in whole living cells using the Magic Red Cathepsin B protease activity Kit (#ICT937, Bio-Rad) according to manufacturer’s instructions. In brief, cells were labelled with the Magic Red substrate and Hoechst (1 μg/ml) for 1 h at 37 °C, and then fixed with 4% formaldehyde in PBS for 10 min at room temperature and mounted on glass slides with Fluoromount-G mounting medium (#00-4958-02, Invitrogen). Samples were imaged using a Leica SP8 inverted laser-scanning confocal microscope using a 100x objective lense. Image analysis and quantification was performed using the Fiji software ^83^, measuring the relative fluorescence intensity per area of at least 50 individual cells per condition per replicate.

### Retention Using Selective Hooks (RUSH) assay

Synchronized trafficking of proteins in the secretory pathway ^56^ was achieved by transfecting WT and GRASP55 KO WI-26 cells with the Str-KDEL-containing PSAP-SBP reporter construct (see also Plasmids). Twenty-four hours post-transfection, cells were treated with 40 μM biotin (#Β4639, Sigma-Aldrich) for different time points as indicated in the figure legends. Next, cells were lysed, supernatants collected and concentrated as described above, and subjected to SDS-PAGE and immunoblotting, or fixed and subjected to indirect immunofluorescence.

### Gene Ontology analysis

Gene Ontology (GO) and pathway enrichment analysis were performed using the Database for Annotation, Visualization and Integrated Discovery (DAVID) tool ^89,90^, as described previously ^47^. In brief, for the analysis of the GRASP55-dependent secretome, proteins whose intensity increases robustly and significantly (log_2_FC > 1, p < 0.05) in the supernatants of GRASP55 KO versus control WI-26 cells were used for GOTERM_CC_FAT, GOTERM_BP_FAT, KEGG_PATHWAY, GOTERM_MF_FAT, and GOTERM_PFAM analyses (Table S1). The human proteome was used as reference list. The data were visualized in a cell plot, generated using DAVID and the associated Flaski apps (https://flaski.age.mpg.de, developed and provided by the MPI-AGE Bioinformatics core facility ^91^) using 9 selected non-redundant significant CC_GO terms.

The full list of proteins detected in the secretome experiment (originally described in ^47^) was used for generating the Volcano plots (gray dots). Proteins whose secretion changes significantly between GRASP55 KO and control cells (p < 0.05) are represented by dark gray dots. Proteins corresponding to the GO term ‘GO:0005764∼lysosome’ are shown as red dots.

### Statistical analysis

Statistical analysis and presentation of quantification data was performed using GraphPad Prism (versions 9 and 10). Information on quantifications for each method is also provided in the respective Methods section. Data in all graphs are shown as mean ± SD. For graphs with only two conditions shown (Fig. 6B, D), significance was calculated using Student’s t-test (unpaired, two-tailed). For all other graphs, significance for the indicated pairwise comparisons was calculated using one-way (Fig. 2-4, S1) or two-way (Fig. 5) ANOVA with *post hoc* Tukey’s multiple comparisons test. Sample sizes (n) and significance values are indicated in figure legends (* p < 0.05, ** p < 0.01, *** p < 0.005, **** p < 0.001).

All findings were reproducible over multiple independent experiments, within a reasonable degree of variability between replicates. For most experiments, at least three independent replicates were performed. The sample size for microscopy experiments (number of individual cells used in quantifications) is provided in the respective figure legends. No statistical method was used to predetermine sample size, which was determined in accordance with standard practices in the field. No data were excluded from the analyses. The experiments were not randomized, and the investigators were not blinded to allocation during experiments and outcome assessment.

## Data availability

The mass spectrometry proteomics data have been deposited to the ProteomeXchange Consortium via the PRIDE ^92^ partner repository with the dataset identifier PXD020331.

## Acknowledgements

We dedicate this manuscript to Dr. Markus Plomann, who sadly passed away in April 2024, during preparation of this manuscript. We thank all members of the Demetriades lab for critical discussions; and the MPI-AGE FACS & Imaging Core Facility for support with confocal microscopy. CD is funded by the European Research Council (ERC) under the European Union’s Horizon 2020 research and innovation programme (grant agreement No 757729), and by the Max Planck Society. JN acknowledges support by the Center for Molecular Medicine Cologne (CMMC) for funding through the Individual Project Funding Program (Project C10) and the Career Advancement Program (Project CAP31). Parts of this work were supported by the Deutsche Forschungsgemeinschaft (DFG, German Research Foundation) through the Research Unit Grant FOR2722 (NU 467/1-1 and DE 3170/1-1; Project No 384170921) to JN and CD, and the DFG Grant PO1539/1-2 (Project No 395238399) to SP. Models in figures created with BioRender.com.

## Author Contributions

Experimental work: JN, assisted by SAF and MT; data analysis: JN, CD; Enzymatic activity assays: MO, SP; project design, conceptualization: JN, CD; supervision: JN, MP, CD; funding acquisition: JN, SP, CD; figure preparation: JN, CD; manuscript draft: JN, CD, with contributions from SP. All authors approved the final version of the manuscript and agree on the content and conclusions.

