## Supplementary material for "GRASP55 Safeguards Proper Lysosome Function by Controlling Sorting of Lysosomal Enzymes at the Golgi": Figures S1-S3

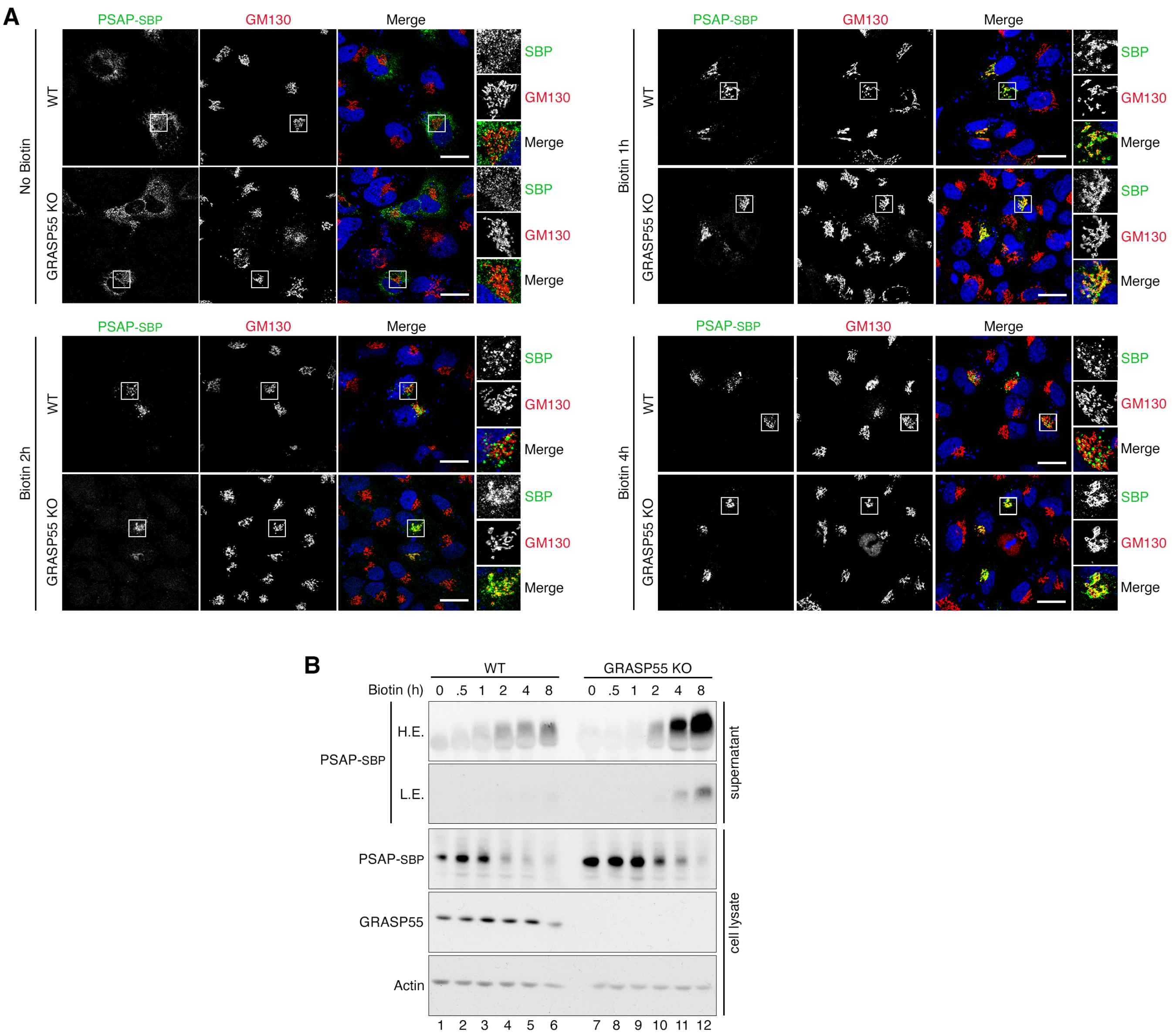


**Figure S1. Missorting of exogenous SBP-tagged PSAP GRASP55 KO cells, detected using a RUSH assay. Related to Figure 1.**

**(A)** Immunofluorescence analysis of WT or GRASP55 KO WI-26 cells transiently expressing streptavidin-binding peptide (SBP)-tagged prosaposin (PSAP) at the indicated time points following biotin addition. GM130 used as a Golgi marker. Nuclei stained with DAPI. Scale bars, 10 µm. n = 3 independent experiments.

**(B)** Representative immunoblot analysis of cell lysates and supernatants from WT or GRASP55 KO WI-26 cells transiently expressing SBP-tagged PSAP at the indicated time points following biotin addition. GRASP55 and Actin immunoblots were used as controls. n = 2 independent experiments.

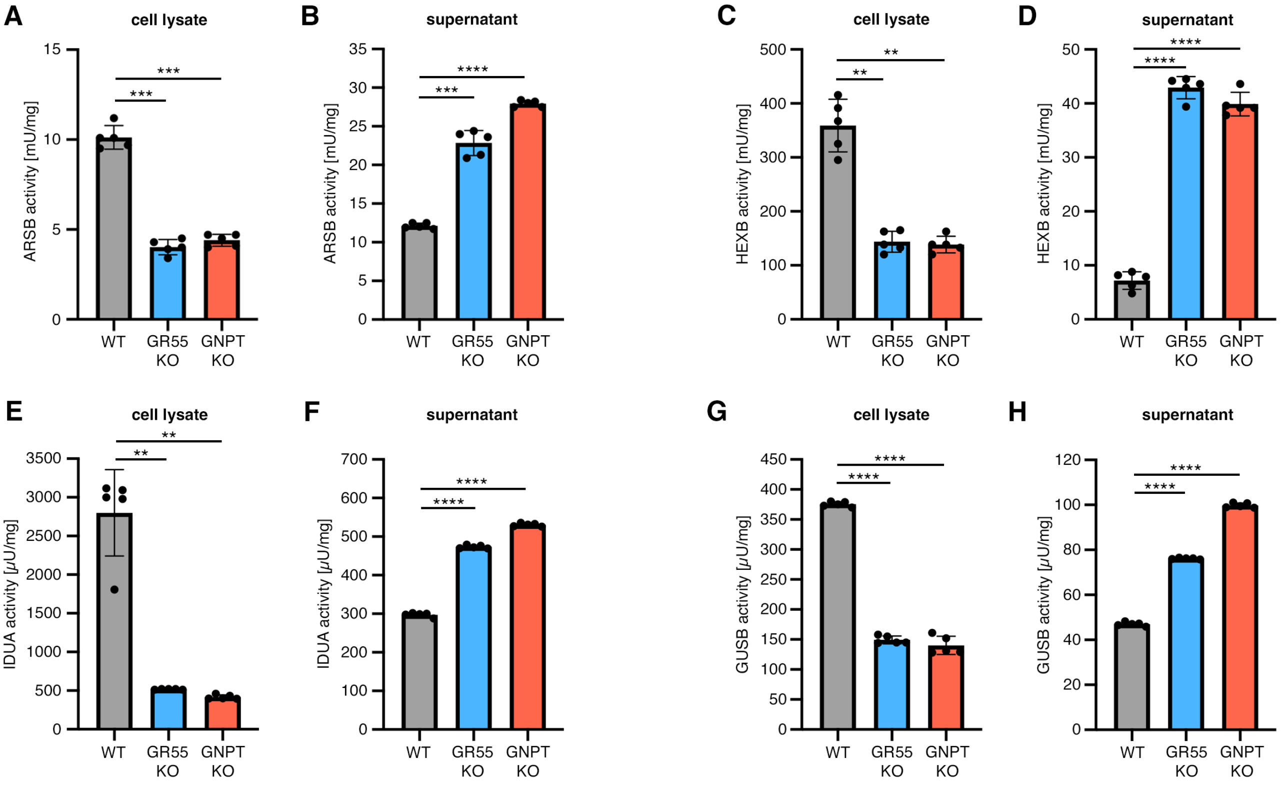


**Figure S2. GRASP55 KO cells exhibit decreased intracellular and elevated extracellular lysosomal enzyme activity, similar to GNPTAB KO cells. Related to Figure 3.**

**(A-B)** The enzymatic activity of arylsulfatase B (ARSB) was assayed in cell lysates (A) or supernatants (B) of wild-type (WT), GRASP55 KO (GR55 KO), or GNPTAB KO (GNPT KO) WI-26 cells (as a cellular model of mucolipidosis type II).

**(C-D)** As in (A-B), but for β-hexosaminidase (HEXB) activity.

**(E-F)** As in (A-B), but for α-L-iduronidase (IDUA) activity.

**(G-H)** As in (A-B), but for β-glucuronidase (GUSB) activity.

n = 5 independent measurements from 4 biological replicates. Data in graphs shown as mean ± SD. ** $p < 0.01$, *** $p < 0.005$, **** $p < 0.001$.

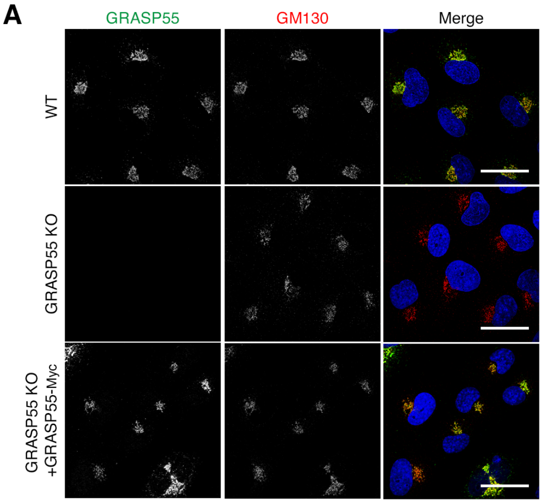


**Figure S3. Stably-expressed myc-tagged GRASP55 localizes at the Golgi, resembling endogenous GRASP55 localization. Related to Figure 4.**

**(A)** Immunofluorescence analysis of WT or GRASP55 KO WI-26 cells, or GRASP55 KO cells stably re-expressing myc-tagged GRASP55 at near-endogenous levels. GM130 used as a Golgi marker. Nuclei stained with DAPI (blue). Scale bar = 10 um. n = 2 independent experiments.
